# Activator protein-1 transactivation of the major immediate early locus is a determinant of cytomegalovirus reactivation from latency

**DOI:** 10.1101/2020.02.06.929281

**Authors:** Benjamin A. Krishna, Amanda B. Wass, Christine M. O’Connor

## Abstract

Human cytomegalovirus (HCMV) is a ubiquitous pathogen that latently infects hematopoietic cells and has the ability to reactivate when triggered by immunological stress. This reactivation causes significant morbidity and mortality in immune-deficient patients, who are unable to control viral dissemination. While a competent immune system helps prevent clinically detectable viremia, a portrait of the factors that induce reactivation following the proper cues remains incomplete. Our understanding of the complex molecular mechanisms underlying latency and reactivation continue to evolve. We previously showed the HCMV-encoded G-protein coupled receptor *US28* is expressed during latency and facilitates latent infection by attenuating the activator protein-1 (AP-1) transcription factor subunit, c-fos, expression and activity. We now show AP-1 is a critical component for HCMV reactivation. Pharmacological inhibition of c-fos significantly attenuates viral reactivation. In agreement, infection with a virus in which we disrupted the proximal AP-1 binding site in the major immediate early (MIE) enhancer results in inefficient reactivation compared to wild type. Concomitantly, AP-1 recruitment to the MIE enhancer is significantly decreased following reactivation of the mutant virus. Further, AP-1 is critical for de-repression of MIE-driven transcripts and downstream early and late genes, while immediate early genes from other loci remain unaffected. Our data also reveal MIE transcripts driven from the MIE promoter, the distal promoter, and the internal promoter, iP2, are dependent upon AP-1 recruitment, while iP1-driven transcripts are AP-1-independent. Collectively, our data demonstrate AP-1 binding to and activation of the MIE enhancer is a key molecular process controlling reactivation from latency.

**Significance Statement:** Human cytomegalovirus (HCMV) is a common pathogen that infects the majority of the population for life. This infection poses little threat in immunologically healthy individuals, but can be fatal in people with compromised immune systems. Our understanding of the mechanisms underlying latency and reactivation remains incomplete. Here, we show the cellular transcription factor, AP-1, is a key to regulating HCMV reactivation. Our findings reveal AP-1 binding to the major immediate early enhancer/promoter is critical for switching this locus from one that is repressed during latency to one that is highly active following reactivation. Our work provides a novel mechanism HCMV exploits to reactivate, highlighting AP-1 as a potential target to prevent HCMV reactivation.

## Introduction

The betaherpesvirus human cytomegalovirus (HCMV) latently infects 50 - 75% of the healthy US population (1). While this infection is, for the most part, asymptomatic in healthy individuals, reactivation from latency causes significant morbidity and mortality in immunocompromised and immunosuppressed individuals (2). Reactivation is triggered by differentiation of latently infected myeloid cells, which when combined with immunodeficiency, results in detectable viremia (3, 4) that is treated with antiviral therapies, such as valganciclovir and foscarnet (5). These antiviral compounds are limited by toxicity and the emergence of drug resistant viral strains. Further, these drugs target viral lytic replication, when disease is already primed to occur (6, 7). Thus, novel approaches aimed at preventing viral reactivation will prove beneficial for immunodeficient patients at risk for HCMV-associated diseases.

Latent infection is defined as the maintenance of viral genomes in the absence of infectious virus production, coupled with the potential to reactivate given the proper extracellular and/or environmental cues (8). A critical determinant in the establishment and maintenance of HCMV latency is repression of the major immediate early promoter (MIEP). The MIEP is powerful promoter, which drives the expression of the major immediate early (MIE) genes, *UL122* and *UL123* that encode IE2 and IE1, respectively. These MIE proteins transactivate other sites of E gene expression, which in turn facilitate the lytic life cycle. As such, the MIE locus is significantly repressed during viral latency (9–12). This silencing is mediated, at least in part, by the recruitment of both repressive transcription factors and chromatin modifiers to the MIE enhancer/promoter region. During reactivation from latency, these repressive factors are overcome, resulting in de-repression of the MIE enhancer/promoter and expression of the MIE-driven genes. This likely requires a multifaceted process that exchanges pro-latency factors for transcription factors and chromatin remodeling favoring the re-expression of IE1 and IE2 (3). It is also clear that additional promoters within the MIE region drive transcription of mRNAs with differing 5’ untranslated region (UTR) lengths, all of which can encode full length IE1 and IE2 proteins. Two of these non-canonical transcripts are derived from promoters within intron A of the MIE region, termed iP1 and iP2 (13). Interestingly, recent work from Collins-McMillen, et al. demonstrated iP2-derived transcripts are the most abundant during both latency and reactivation. Furthermore, transactivation of both iP1 and iP2 are required for efficient viral reactivation in primary CD34^+^ hematopoietic progenitor cells (HPCs) (14), though the regulation of these newly identified promoters remains unknown. These findings highlight the complexity of the transcriptional control of the MIE locus.

Work from many investigators implicates multiple factors in de-repressing the MIE locus during reactivation. These include: src family kinase activation of extracellular signal-regulated kinase/mitogen-activated protein kinase (ERK/MAPK; ref. (15)), cyclic AMP (cAMP) activation (16), cAMP response element-binding protein (CREB) phosphorylation (17), epidermal growth factor receptor (EGFR) activation of phosphoinositide 3-kinase (PI3K; ref. (18)), facilitates chromatin transcription (FACT)-mediated transactivation (19), mammalian target of rapamycin (mTOR)-modulated activation of KRAB-associated protein-1 (KAP1; ref. (20)), and protein kinase A (PKA)-CREB-TORC2 transactivation (21). Although these results summarize findings from a variety of latency model systems, they underscore the idea that reactivation likely requires concomitant signals from multiple pathways and factors to de-repress the MIE locus, triggering epigenetic changes as well as the recruitment of transcription factors that transactivate this region.

We recently showed *US28*, an HCMV encoded G-protein coupled receptor (GPCR), attenuates cellular fos (c-fos) signaling to maintain latency (22). Dimerization of c-fos and c-jun form the transcription factor, activator protein-1 (AP-1), which binds and transactivates the MIEP in luciferase reporter assays (23). US28 protein (pUS28) suppression of phosphorylated c-fos prevents AP-1 from binding the MIE enhancer and activating the MIE locus during latent infection of Kasumi-3 cells (a CD34^+^ cell line that supports latency and viral reactivation) and primary *ex vivo* CD34^+^ hematopoietic progenitor cells (HPCs) (22). The MIE enhancer region contains two AP-1 binding sites: the promoter proximal binding site has a canonical AP-1 binding motif, while the promoter distal binding site has a non-consensus sequence (23–25). Disruption of both binding sites together does not impact lytic replication in fibroblasts or epithelial cells, nor does it affect virulence in a chimeric murine CMV infection model (23), suggesting AP-1 recruitment to the MIEP is dispensable for lytic infection.

Our previous data revealed suppressing AP-1 activity is necessary for maintaining HCMV latency (22), and thus we hypothesized AP-1 recruitment to the MIEP is important for efficient reactivation from latency in hematopoietic cells. Herein, we show pharmacological inhibition of AP-1 in the presence of reactivation stimuli decreases reactivation efficiency. Additionally, mutation of the promoter proximal AP-1 binding site attenuates AP-1 recruitment to the MIE enhancer and impairs reactivation following stimuli. Finally, our data reveal AP-1 recruitment to the promoter proximal site in the MIE enhancer during reactivation is required to stimulate the expression of MIE-driven genes originating from the canonical MIEP, iP2, and the distal promoter (dP), but not other IE genes or iP1-driven transcripts. Taken together, our results show AP-1 transactivation of the MIE enhancer/promoter is an important factor for efficient HCMV reactivation from latency.

## Results

### Pharmacological inhibition of c-fos suppresses viral reactivation in Kasumi-3 cells

We previously showed HCMV suppresses AP-1 activation to maintain latency (22), which led us to posit the requirement for this transcription factor’s activation during reactivation from latency. To test the impact AP-1 signaling has on reactivation, we infected Kasumi-3 cells with TB40/E*mCherry* (WT) and cultured the cells in conditions favoring latency for 7 days (d). Cultures were then treated with vehicle (dimethylsulfoxide, DMSO) to maintain latency or tetradecanoyl phorbol acetate (TPA) to induce reactivation for an additional 2 d in the presence or absence of the c-fos inhibitor, T-5224, at a concentration we previously showed does not impact cell viability (22). We assessed the ability of each infection to reactivate by co-culturing the infected Kasumi-3 cells with naïve fibroblasts and quantified infectious virus production by Extreme Limiting Dilution Analysis (ELDA). As expected, TPA treatment of infected Kasumi-3 cells induced reactivation, as measured by an increase in virus production (26). However, treatment with T-5224 attenuated virus production in TPA-treated cells (Fig. 1A). We next confirmed this finding using cord blood-derived, *ex vivo* cultured CD34^+^ HPCs, a natural site of HCMV latency. We infected these primary cells with WT virus for 7 d under conditions favoring latency, after which we cultured a portion of the infected cells in media that promotes reactivation (27) either with or without T-5224. Similar to Kasumi-3 cells, our findings reveal treatment with T-5224 reduced virus production in CD34^+^ HPCs treated with reactivation stimuli (Fig. 1B). Together, these results indicate AP-1 activation is important for efficient reactivation from latency.

**Figure 1.**
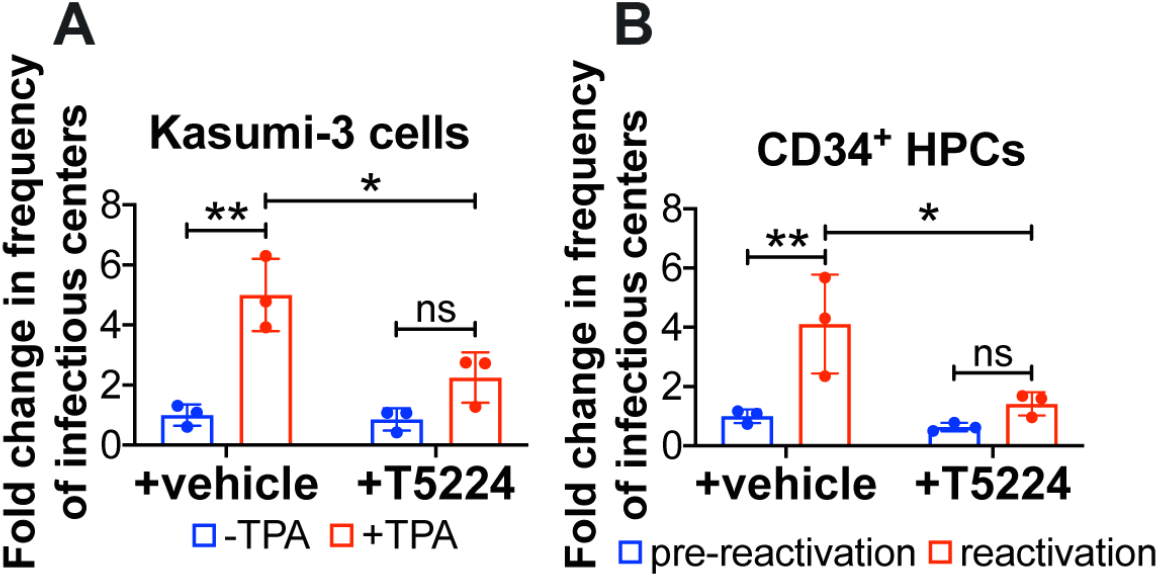
Selective, pharmacological inhibition of c-fos/AP-1 suppresses viral reactivation. **(A)** Kasumi-3 cells (moi = 1.0) or **(B)** CD34^+^ HPCs (moi = 2.0) were infected for 7 d under latent conditions with WT. Infectious particles were quantified by Extreme Limiting Dilution Analysis (ELDA) with **(A)** DMSO (-TPA, blue bars) or with TPA (+TPA, red bars) or **(B)** hLTCM (pre-reactivation, blue bars) or reactivation media (red bars), in the presence of vehicle (DMSO) or the c-fos inhibitor, T5224 (20 nM). Frequency of infectious centers is plotted as fold-change relative to WT +vehicle under latent conditions. Each data point (circles) is the mean of 3 technical replicates. Error bars indicate standard deviation of three biological replicates, each of which is shown in SI Appendix, Fig. S1. Statistical significance was calculated using two-way ANOVA followed by Tukey’s post-hoc analysis. \**p*<0.05, \*\**p*<0.01, ns = not significant

### Mutation of the promoter proximal AP-1 binding site in the MIE enhancer impairs viral reactivation in hematopoietic cells

Based on our findings above, we next asked if AP-1 recruitment to the MIE enhancer aids in de-repressing the MIE locus during HCMV reactivation. To begin to investigate the contribution of AP-1 to this process, we generated a recombinant virus in the bacterial artificial chromosome (BAC)-derived, clinical isolate TB40/E (28), which we previously engineered to express mCherry (TB40/E*mCherry*) (29). We disrupted the promoter proximal AP-1 binding site to generate TB40/E*mCherry*-proximal-AP-1*mutant* (AP-1*mut_p_*). Additionally, to ensure we did not introduce off-site mutations during the recombineering process, we restored the mutated MIE enhancer/promoter region to its wild type sequence to generate the repair virus, TB40/E*mCherry*-proximal-AP-1*repair* (AP-1*rep_p_*) (SI Appendix, Fig. S2). Consistent with previous findings (23), we confirmed AP-1*mut_p_* and AP-1*rep_p_* display WT growth phenotypes in fibroblasts (SI Appendix, Fig. S3A). Similarly, both the mutant and repair viruses displayed WT growth following lytic infection of epithelial cells (SI Appendix, Fig. S3B). Together, these results suggest AP-1 recruitment to the promoter proximal site is not required for efficient lytic replication.

To determine the contribution of the AP-1 promoter proximal binding site during reactivation from latency, we infected Kasumi-3 cells with WT, AP-1*mut_p_*, or AP-1*rep_p_* for 7 d under conditions favoring latency, after which we performed ELDA in the presence of vehicle or TPA treatment. We found AP-1*mut_p_*-infected Kasumi-3 cells failed to efficiently reactivate to WT or AP-1*rep_p_* levels (Fig. 2A). Importantly, AP-1*mut_p_*-infected cells maintain comparable levels of the viral genome compared to WT-or AP-1*rep_p_*-infected cultures (SI Appendix, Fig. S5), suggesting these infected cultures harbor latent virus. Similarly, the latency-associated gene, *UL138*, was highly expressed relative to *UL123* in all infected cultures, consistent with latent infection (SI Appendix, Fig. S6). Treatment of each infected culture with TPA resulted in a decrease in the ratio of *UL138*/*UL123* (SI Appendix, Fig. S6), consistent with viral reactivation (22, 26, 30). The ratio of *UL138*:*UL123* neared 1 in the WT- and AP-1*rep_p_*-infected cells following TPA treatment, which was significantly lower than the ratio of these transcripts in the AP-1*mut_p_*-infected counterpart cultures (SI Appendix, Fig. S6), suggesting inefficient reactivation of the mutant virus. Consistent with these observations, AP-1*mut_p_*-infected primary CD34^+^ HPCs produces significantly less infectious virus than either WT or AP-1*rep_p_* when stimulated to reactivate (Fig. 2B). Together, these data suggest AP-1*mut_p_*-infected hematopoietic cells undergo latency, but fail to efficiently reactivate to WT levels following stimuli.

**Figure 2.**
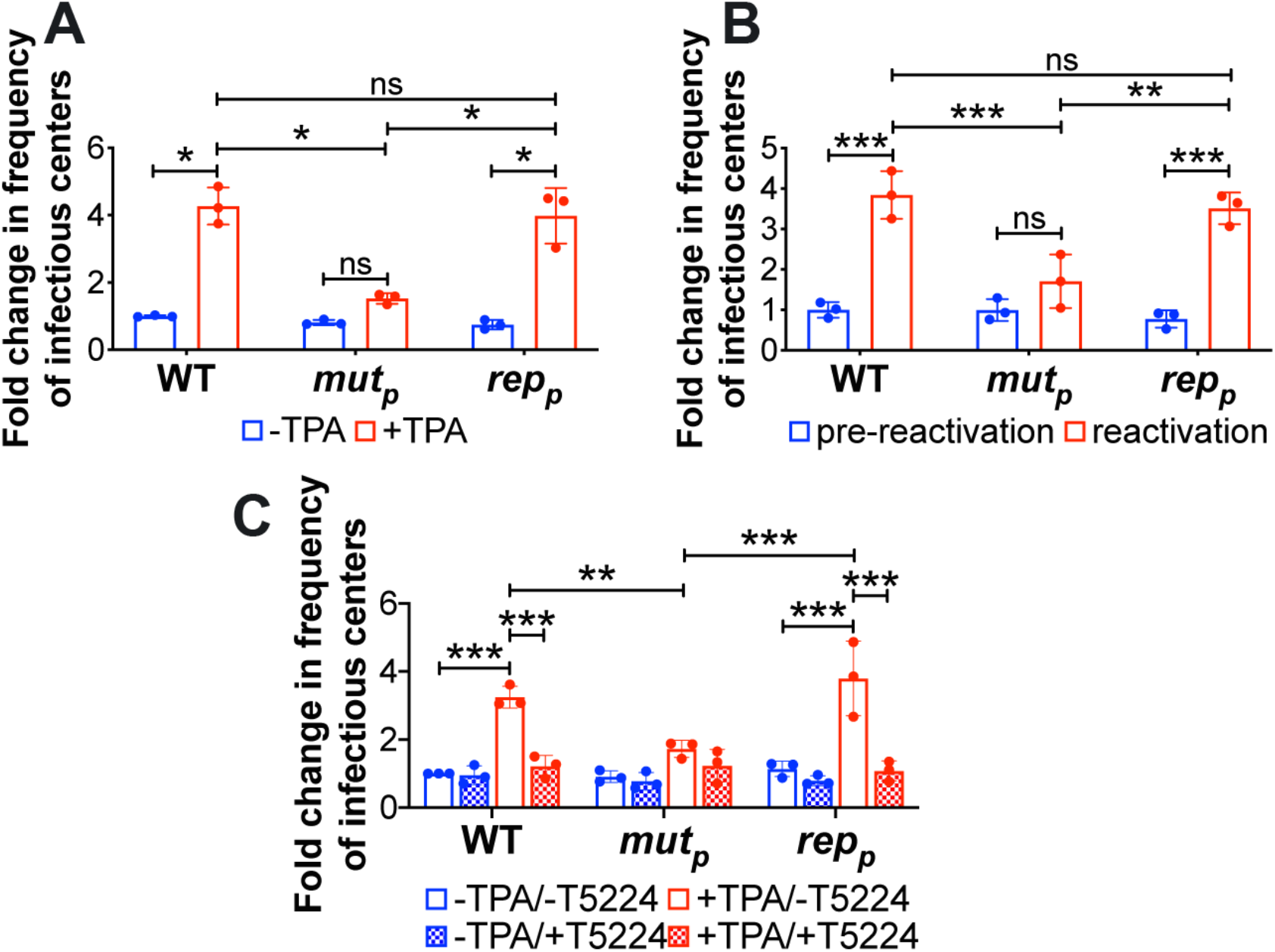
Mutation of the promoter proximal AP-1 binding site in the MIE enhancer decreases efficient viral reactivation in Kasumi-3 and primary CD34^+^ cells. **(A,C)** Kasumi-3 cells (moi = 1.0) or **(B)** cord blood-derived CD34^+^ HPCs (moi = 2.0) were infected with the indicated viruses. At 7 dpi, half of each infected population was cultured for an additional 2 d **(A,C)** with vehicle (DMSO; −TPA, blue bars) or TPA (+TPA, red bars) or **(B)**with hLTCM (pre-reactivation, blue bars) or reactivation (red bars) media. **(A,B)** Cells were then co-cultured with naïve NuFF-1 cells to quantify the frequency of infectious centers by ELDA. **(C)** Cells were co-cultured with naïve NuFF-1 cells in the presence of vehicle (DMSO; -T5224, open bars) or the fos inhibitor, T5224 (checked bars), and the frequency of infectious particles was quantified by ELDA. **(A-C)** Each data point (circles) is the mean of 3 technical replicates. Error bars indicate standard deviation of three biological replicates, each of which is shown in SI Appendix, Fig. S4. Data is presented as fold-change relative to WT **(A)** −TPA, **(B)** pre-reactivation, or **(C)** −TPA/−T5224. The statistical significance was calculated using two-way ANOVA followed by Tukey’s post-hoc analysis. *p<0.05, \*\**p*<0.01, \*\*\**p*<0.001, and ns = not significant. Abbreviations: AP-1*mut_p_* (*mut_p_*), AP-1*rep_p_* (*rep_p_*).

Finally, to distinguish between the relative contributions of AP-1 activity and the requirement for the AP-1 promoter proximal binding site during reactivation from latency, we evaluated viral reactivation in the presence or absence of the c-fos inhibitor, T-5224. To this end, we infected Kasumi-3 cells with WT, AP-1*mut_p_*, or AP-1*rep_p_* for 7 d under conditions favoring latency. We then quantified infectious virus production by ELDA in the presence or absence of TPA and/or T-5224. In the presence of T-5224, WT- and AP-1*rep_p_*-infected cells fail to efficiently reactivate virus, resulting in virion production similar to AP-1*mut_p_*-infected cells in the absence of T-5224 treatment (Fig. 2C). Further, T-5224 did not further impair reactivation of the AP-1*mut_p_* virus. Taken together, these results demonstrate the importance of AP-1 activation and the requirement for the promoter proximal site of the MIE enhancer in promoting HCMV reactivation.

### Recruitment of the AP-1 transcription factor to the MIE enhancer is significantly diminished following reactivation stimulus of AP-1*mut_p_*-infected Kasumi-3 cells

To test our hypothesis that AP-1 binds to the MIE enhancer, we first evaluated the recruitment of the AP-1 subunits, c-fos and c-jun, to the enhancer region. We latently infected Kasumi-3 cells with WT, AP-1*mut_p_*, or AP-1*rep_p_* for 7 d and then treated a portion of the infected cultures for an additional 2 d with TPA treatment to induce reactivation. We then harvested the cultures and performed chromatin immunoprecipitation (ChIP) assays to measure binding of c-fos and c-jun to the MIE enhancer. Our results show both c-fos and c-jun bind the MIE enhancer following TPA treatment of WT, latently infected Kasumi-3 cells, although disruption of the promoter proximal AP-1 site reduced recruitment of each AP-1 subunit (Fig. 3). Importantly, AP-1*mut_p_* infection does not impact c-fos or c-jun transcript levels, which are comparable to those in WT- and AP-1*rep_p_*-infected cells both prior to and following TPA treatment (SI Appendix, Fig. S7). Further, consistent with previous findings (22, 31), c-fos and c-jun mRNA is attenuated during latent infection relative to conditions favoring reactivation (SI Appendix, Fig. S7). While both c-fos and c-jun recruitment is significantly attenuated in the AP-1*mut_p_*-infected Kasumi-3 cells, our data reveal neither AP-1 subunit is completely devoid of binding. This suggests AP-1 also binds the promoter distal, non-consensus AP-1 binding site in the MIE enhancer region. To address this, we generated the AP-1 promoter distal mutant, termed AP-1*mut_d_*, and its match repair virus, AP-1*rep_d_*. We also generated a viral mutant in which we mutated both proximal and distal AP-1 binding sites in a single background, termed AP-1*mut_pd_*, as well as a repair virus, in which we returned both mutated sequences to wild type, AP-1*rep_pd_* (SI Appendix, Fig. S2). Each of these viruses replicated to wild type titers in primary fibroblasts (SI Appendix, Fig. S8A), indicating AP-1 binding to the MIE enhancer is not required for efficient lytic replication. However, viral reactivation was significantly impaired in AP-1*mut_pd_*-infected Kasumi-3 cells treated with TPA, which was not significantly different when compared to cells infected with AP-1*mut_p_*, whereas AP-1*mut_d_*-infected Kasumi-3 cells reactivated to WT levels following TPA treatment (SI Appendix, Fig. 8B). Further, both c-fos and c-jun subunits of AP-1 bound the MIE enhancer in AP-1*mut_d_*-infected, but not AP-1*mut_pd_*-infected Kasumi-3 cells treated with TPA (SI Appendix, Fig. 8C and D). Our data collectively indicate the promoter distal AP-1 binding site does not play a significant role in reactivation from latency, whereas deletion of the promoter proximal site alone significantly impacts efficient reactivation following stimuli (Fig. 2A and B, SI Appendix, Fig. 8B), despite detectable promoter distal site binding of AP-1 (Fig. 3, SI Appendix, Fig. 8C and D).

**Figure 3.**
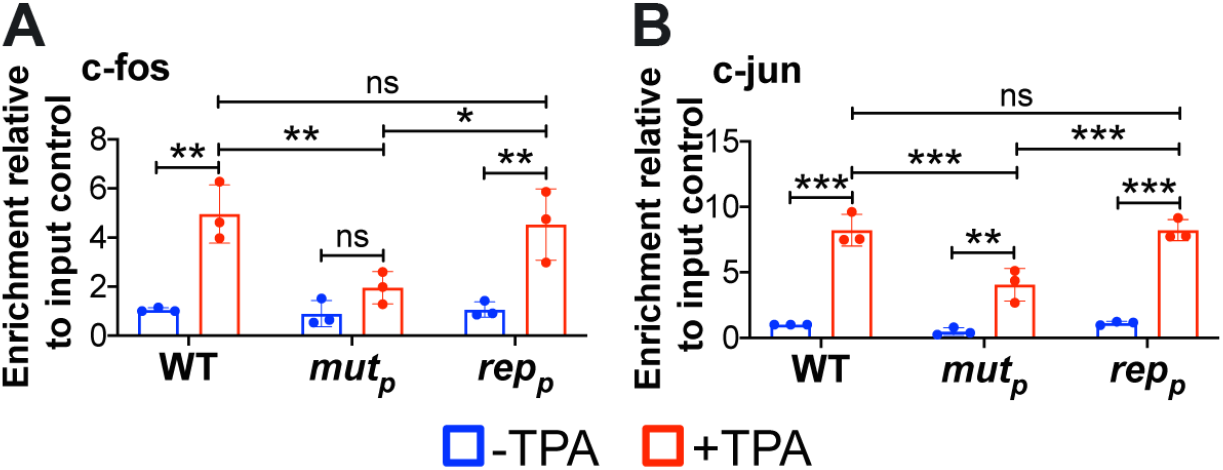
AP-1 *mut_p_* infection of Kasumi-3 cells results in a significant reduction of AP-1 transcription factor binding to the MIE enhancer in the presence of reactivation stimuli. Kasumi-3 cells were infected (moi = 1.0) with the indicated viruses. At 7 dpi, half of each infected population was cultured for an additional 2 d with vehicle (DMSO; −TPA, blue bars) or TPA (+TPA, red bars). AP-1 transcription factor binding to the MIEP was quantified by ChIP using **(A)** α-fos or **(B)** α-jun antibodies. Co-precipitated MIEP was quantified by qPCR, and data are shown as fold change relative to input. Each data point (circles) is the mean of 3 technical replicates. Error bars indicate standard deviation of three biological replicates, and the statistical significance was calculated using two-way ANOVA followed by Tukey’s post-hoc analysis. \**p*<0.05, \*\**p*<0.01, \*\*\**p*<0.001, and ns = not significant. Abbreviations: AP-1*mut_p_* (*mut_p_*), AP-1*rep_p_* (*rep_p_*).

### AP-1 recruitment to the MIE enhancer during reactivation initiates MIE-driven gene expression, but not the transcription of other IE genes

Together, our data reveal AP-1 recruitment to the promoter proximal site in the MIE enhancer is an important determinant for efficient viral reactivation. Thus, we hypothesized this transcription factor is critical for de-repression of *UL122* and *UL123* expression. In line with this, we posited the recruitment of AP-1 to the MIE enhancer would not impact other IE genes. To this end, we latently infected Kasumi-3 cells with WT, AP-1*mut_p_*, or AP-1*rep_p_* for 7 d, after which we treated a portion of the cells with TPA for an additional 2 d to stimulate reactivation. We then harvested total RNA from the infected cultures and analyzed HCMV gene expression by RT-qPCR. We assessed the MIE-driven transcripts *UL122* and *UL123*, representative non-MIE-derived IE genes (*UL36*, *UL37*), as well as a representative early (E; *UL44*) and late (L; *UL99*) transcripts. AP-1*mut_p_*-infected Kasumi-3 cells treated with TPA displayed attenuated *UL122* and *UL123* transcription compared to the WT-or AP-1*rep_p_*-infected cultures (Fig. 4A and B). As expected, this decreased E (*UL44*, Fig. 4E) and L (*UL99*, Fig. 4F) transcript abundance, whose transactivation is dependent upon MIE-driven transcription and their subsequent translation (32). However, transcription of *UL36* and *UL37*, whose expression is not regulated by the MIE enhancer, was unaffected by the mutation of the promoter proximal AP-1 site (Fig. 4C and D). Taken together, these data suggest AP-1 recruitment to the promoter proximal site within the MIE enhancer is required for the expression of MIE-driven transcripts and efficient viral reactivation.

**Figure 4.**
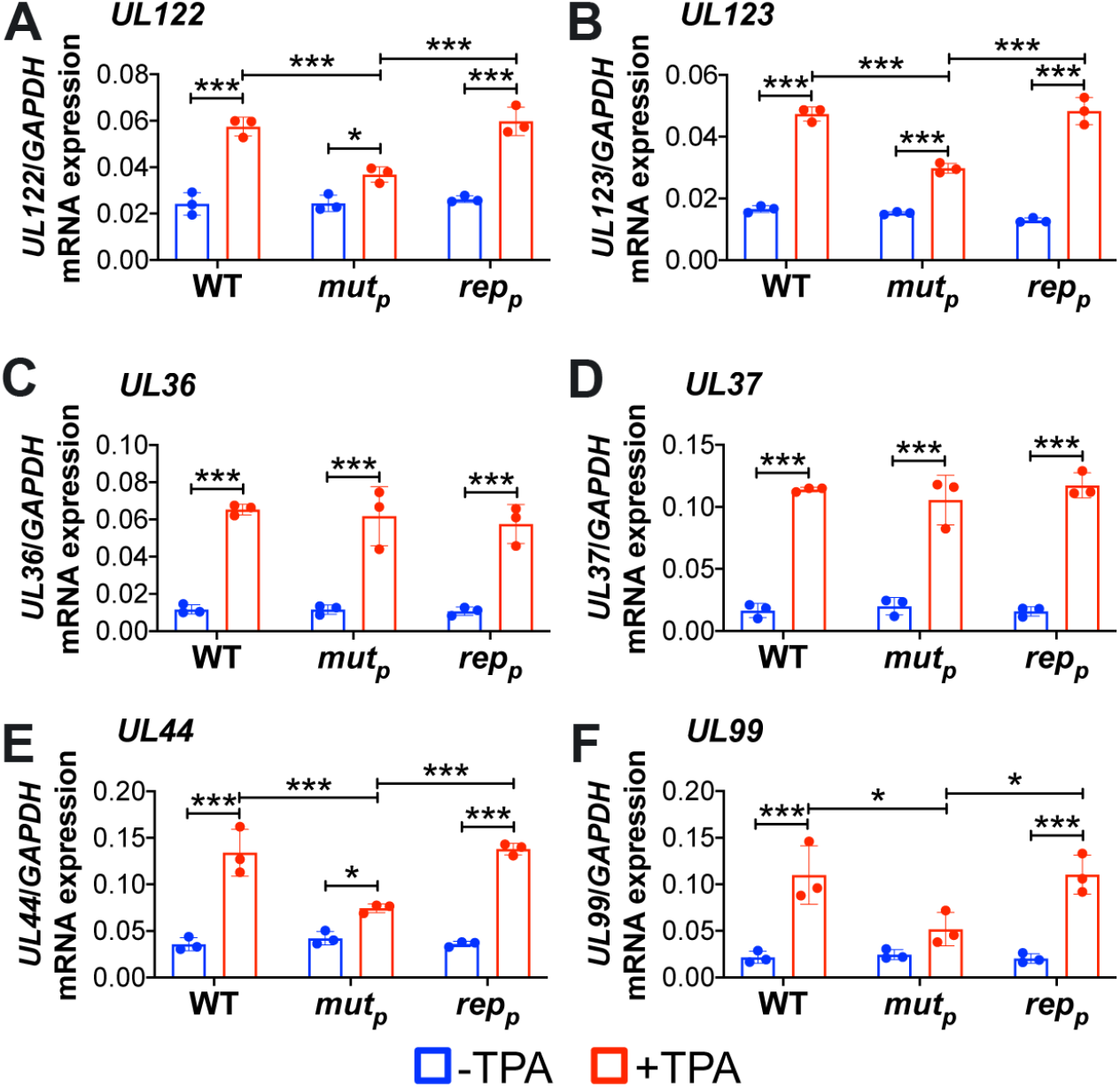
AP-1 *mut_p_* infection of Kasumi-3 cells impacts MIEP-driven gene expression, but not the transcription of other IE genes. Kasumi-3 cells were latently infected (moi = 1.0) with the indicated viruses. At 7 dpi, half of each infected population was cultured for an additional 2 d with vehicle (DMSO) to maintain latency (-TPA, blue bars) or with TPA to induce reactivation (red bars). Representative **(A,B)** MIE, **(C,D)** IE, **(E)** E, and **(F)** L genes were assessed by RTqPCR relative to cellular *GAPDH* and are plotted as arbitrary units. Each data point (circles) is the mean of 3 technical replicates. Error bars indicate standard deviation of three biological replicates. The statistical significance was calculated using two-way ANOVA followed by Tukey’s post-hoc analysis. \**p*<0.05, \*\*\**p*<0.001. Abbreviations: AP-1*mut_p_* (*mut_p_*), AP-1*rep_p_* (*rep_p_*).

Thus far, our findings indicate disruption of promoter proximal AP-1 binding to the MIE enhancer impairs de-repression of MIE transcript expression. However, IE1 and IE2 are translated from multiple transcripts originating from the canonical MIEP, as well as alternative transcription start sites (TSSs) within the MIE region. While these alternative transcripts differ in their 5’UTR, each encodes full-length IE1 or IE2 protein (13). More recently, Collins-McMillen et al. found two of these alternative transcripts derived from promoters within intron A, iP1 and iP2, are important for reactivation from latency (14). However, the contribution of transcription factor binding on the activity of these alternative promoters during reactivation is unknown. To this end, we infected Kasumi-3 cells as above and analyzed expression of MIE mRNAs by RT-qPCR, using primers specific for the transcripts derived from the dP, canonical MIEP, or the internal promoters, iP1 and iP2 (Fig. 5A). Following TPA-induced reactivation of WT-infected cultures, transcripts originating from each promoter increased (Fig. 5B-E), concomitant with *UL122* (Fig. 4A) and *UL123* (Fig. 4B) transcription. These findings are consistent with previous findings using the THP-1 culture system (14). However, AP-1*mut_p_*-infected Kasumi-3 cells displayed impaired MIEP-, iP2-, and dP-derived transcripts compared to cultures infected with either WT or AP-1*rep_p_* (Fig. 5B, D, and E, respectively). In contrast to these MIE-derived transcripts, mRNA driven from the iP1 promoter was expressed to similar levels following reactivation when compared to WT-or AP-1*rep_p_*-infected cultures (Fig. 5C), suggesting iP1-driven transcription is independent of AP-1 binding to the promoter proximal site in the MIE enhancer. Taken together, these data suggest AP-1 binding to the MIE enhancer facilitates the expression of MIEP-, iP2-, and dP-derived transcripts, while iP1-driven transcription remains independent of AP-1 recruitment to this region.

**Figure 5.**
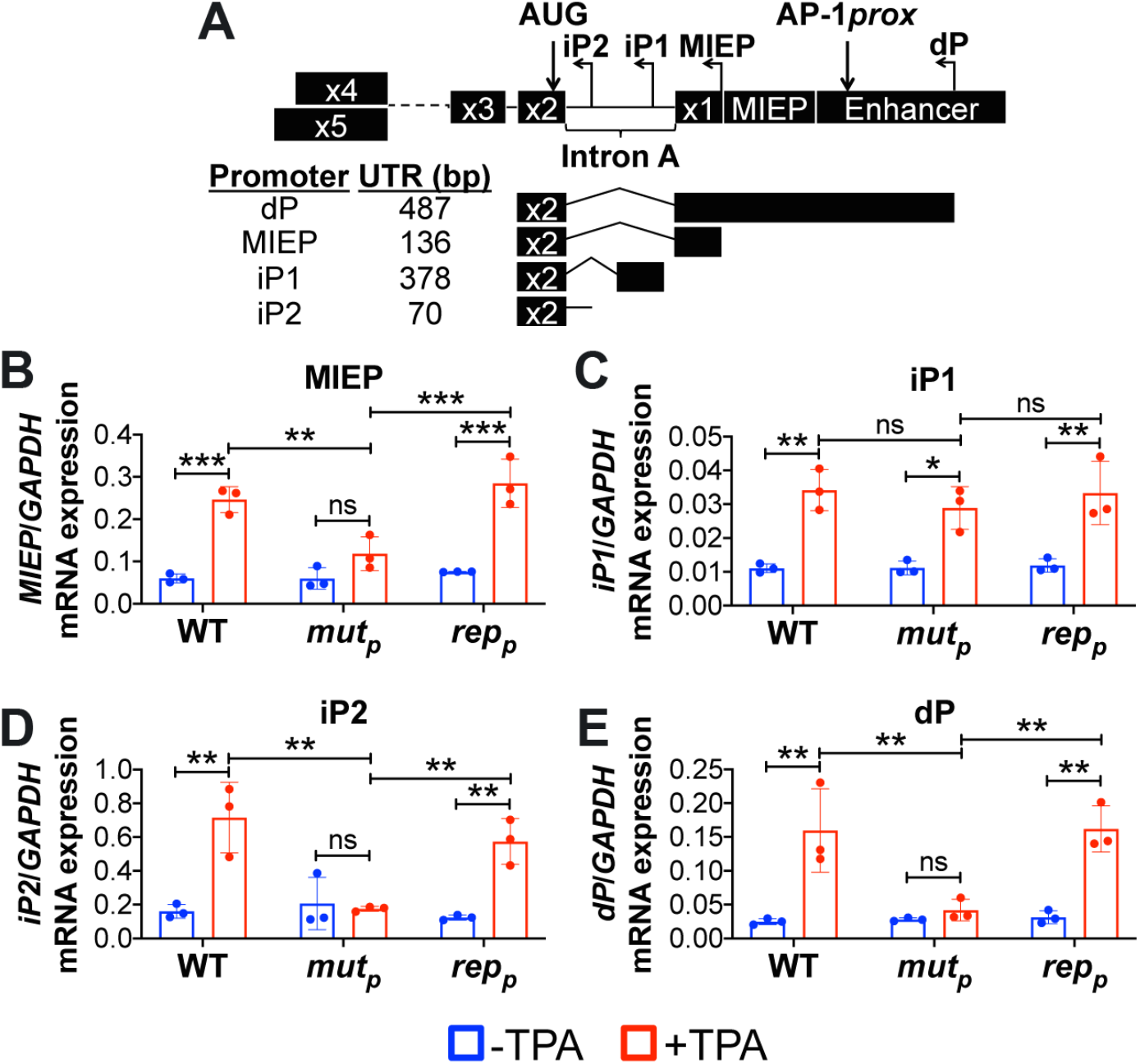
Transcripts driven from the canonical MIEP, iP2, and dP require AP-1 binding following reactivation. Kasumi-3 cells were infected (moi = 1.0) with the indicated viruses. At 7 dpi, half of each infected population was cultured for an additional 2 d with vehicle (DMSO; - TPA, blue bars) or TPA (+TPA, red bars). Transcripts derived from the promoters within the MIE region (not to scale) depicted in **(A)** were assessed: **(B)** MIEP; intronic promoters, **(C)** iP1 and **(D)** iP2; and **(E)** the distal promoter (dP). **(B-E)** Expression of each transcript was measured by RTqPCR and is shown relative to cellular *GAPDH,* plotted as arbitrary units. Each data point (circles) is the mean of 3 technical replicates, and the error bars indicate standard deviation of three biological replicates. Statistical significance was calculated using two-way ANOVA followed by Tukey’s post-hoc analysis. \**p*<0.05, \*\**p*<0.01, \*\*\**p*<0.001, and ns = not significant. Abbreviations: AP-1*mut_p_* (*mut_p_*), AP-1*rep_p_* (*rep_p_*).

## Discussion

HCMV reactivation is a multifaceted process, requiring de-repression of *UL122* and *UL123* transcription, changes in transcription factor binding, as well as changes in chromatin structure to promote transcription (3). In this study, we found AP-1 binding to the MIE enhancer is critical for the re-expression of both *UL122* and *UL123*, as well as HCMV reactivation. Pharmacological inhibition of c-fos, a component of AP-1, significantly reduced the reactivation efficiency of WT-infected hematopoietic cells (Fig. 1). Similarly, disruption of the promoter proximal AP-1 binding site within the MIE enhancer attenuated HCMV reactivation (Fig. 2). Our data demonstrate AP-1 is recruited to the MIE enhancer following reactivation of WT virus, and mutation of the promoter proximal AP-1 binding site significantly decreases AP-1 binding (Fig. 3). Further, while the MIE enhancer has two AP-1 binding sites, our data emphasize the promoter proximal site is critical for viral reactivation (Fig. 2, 3, SI Appendix Fig. S8). AP-1 recruitment to the MIE enhancer aids in the transactivation of the MIE-driven IE genes, *UL122* and *UL123*, while IE genes not derived from the MIE region, including *UL36* and *UL37*, remain unaffected (Fig. 4). Importantly, AP-1 binding to its promoter proximal site in the MIE enhancer activates transcription from the canonical MIEP, as well as the alternative iP2 and dP promoters, while iP1 activity is AP-1-independent (Fig. 5). Together, our results reveal a mechanism by which the AP-1 transcription factor binds the MIE enhancer to activate multiple promoters in the MIE locus, thereby contributing to successful reactivation of HCMV from latency.

While viral reactivation is likely multifactorial, it is clear c-fos and c-jun are critical components of this process. Our previous findings showed pUS28 attenuates c-fos expression and activation, concomitant with reduced AP-1 recruitment to the MIE enhancer during latency (22). Additionally, we have shown c-jun expression is significantly suppressed during HCMV latency (31), although this is not contingent upon pUS28 expression or function (22). These findings suggest HCMV has adapted independent mechanisms to attenuate the two subunits comprising AP-1 to prevent its binding to the MIE enhancer during latency.

Our findings also suggest HCMV evolved sites for AP-1 recruitment to the MIE enhancer region for viral reactivation, although this association is not required for efficient lytic replication. This is not unprecedented, as similar mechanisms were described for the CRE response elements (17, 33). Consistent with this, several groups have shown AP-1 binds to and activates the MIE enhancer/promoter in *in vitro* expression assays (23, 33, 34). In agreement with our results herein, AP-1 is recruited to the MIE enhancer in a murine model of allograft-induced murine CMV (MCMV) reactivation (35). Furthermore, work from Isern, et al, revealed disruption of both AP-1 binding sites within the MIE enhancer region had no effect on HCMV lytic replication in fibroblast and epithelial cells (23), consistent with our current findings. Additionally, Isern, et al. generated a chimeric MCMV, replacing the native murine MIE enhancer with the HCMV enhancer region, and then disrupted both AP-1 binding sites. Using this chimeric recombinant in either lung or spleen explants of neonatal mice, they found no difference in virus reactivation relative to infection with a chimeric virus containing wild type AP-1 binding sites (23). While this work revealed no role for the promoter proximal AP-1 binding site, there are significant differences in the experimental models we employed herein and the chimeric MCMV-infected mouse models used in the previously published study (23).

In addition to HCMV, other herpesviruses modulate AP-1 during infection. The Epstein Barr virus (EBV) protein, BGLF2, and the Kaposi’s sarcoma-associated herpesvirus (KSHV)-encoded ORF45 both activate the AP-1 signaling pathway to promote viral gene expression, replication, and survival (36, 37). EBV also encodes an AP-1 homolog, BZLF1, which, like AP-1, supports resting B cell proliferation and binds methylated EBV promoters critical for reactivation (38). Murine gammaherpesvirus 68 (MHV68) expresses the microRNA mghv-miR-M1-1, which downregulates c-jun and subsequent AP-1 activation, resulting in the suppression of viral lytic replication (39). Together, these results show controlling AP-1 is a common strategy among herpesviruses, suggesting targeting this transcription factor could prove beneficial for suppressing reactivation and subsequent disease associated with a variety of herpesviruses.

Recently published findings from Collins-McMillen, et al. detailed the requirement for iP1- and iP2-derived transcripts for viral reactivation (14). Indeed, consistent with their findings, we also found the most abundant transcripts are derived from the iP2 alternative promoter, both before and after reactivation, although MIEP-derived transcripts were more abundant than those transcribed from iP1 in our system (Fig. 5). While our overall goal was not to discern differences in abundances, this difference could be due to the systems used: we measured these transcripts in Kasumi-3 cells, while Collins-McMillen determined transcript abundance over time in THP-1 cells (14). Nonetheless, our findings build upon these prior observations and begin to unravel the likely complex web of transcription factors that regulate the iP1 and iP2 promoters. We found AP-1 recruitment to the promoter proximal site within the MIE enhancer drives transcription from the MIEP, iP2, and dP (Fig. 5B, D, and E, respectively). However, transcription from iP1 is independent of AP-1 recruitment to the promoter proximal site within the MIE enhancer, as the AP-1*mut_p_*-infected cells express iP1-driven transcripts to levels similar to those we observed in WT-infected cells (Fig. 5C). This finding suggests iP1-driven transcription alone is not sufficient to drive reactivation, as AP-1*mut_p_*-infected cells fail to efficiently produce infectious virus following the addition of reactivation stimuli (Fig. 2). This result also suggests additional transcription factors or chromatin remodeling proteins may alternatively control iP1 activity, such as FOXO3A (personal communication, Nat Moorman), further highlighting the complexity in the regulation of transcription in MIE locus.

How is AP-1 activated in response to cues that reactivate HCMV? Elucidating the precise biological mechanism could indeed reveal potential therapeutic targets one could exploit to prevent viral reactivation. Two pathways that activate AP-1, Src-ERK (40) and PI3K/Akt (41), are already implicated in HCMV reactivation (15, 18) and are therefore potential candidates for further investigation. As a potent signaling molecule, US28 is an attractive, latently expressed viral protein capable of modulating cellular signaling pathways such as these. While our previous data showed US28 attenuates c-fos expression and activity as well as AP-1 binding to the MIE enhancer during latency (22), it remains unknown how US28 signaling is modified to favor reactivation rather than latent conditions. It is attractive to hypothesize an additional viral or cellular protein is triggered to alter US28’s behavior following reactivation stimuli, though additional work is needed to realize such mechanistic nuances. It is also possible AP-1 recruits other factors, such as histone acetyltransferases (HATs) (42) to facilitate chromatin remodeling and/or other transcription factors (e.g. CREB and NFκB), all of which would contribute to de-repression of MIE promoters (3). In reporter assays, pp71 transactivates promoters containing AP-1 or CREB in human fibroblasts (43). While pp71, in conjunction with cellular DAXX and ATRX, aids in silencing the MIE enhancer/promoter via repressive heterochromatic marks and histone deposition (44), whether pp71 transactivates the MIE enhancer/promoter during reactivation via AP-1 and or CREB, remains elusive. Finally, c-fos and c-jun functional activity contribute to myeloid differentiation and function in response to pro-inflammatory stimuli (45). These possibilities present numerous avenues for future research aimed at understanding the underlying biological mechanisms regulating AP-1 activation and its transactivation of the MIE enhancer/promoter region.

Our findings detailed herein reveal AP-1 activation is critical for HCMV reactivation. AP-1 recruitment to its promoter proximal site within the MIE enhancer activates multiple MIE promoters, including the dP, MIEP, and iP2, to drive MIE transcription, which overall facilitates reactivation. Herein, we provide a novel mechanism underlying MIE de-repression during HCMV reactivation, further illuminating the molecular details essential for understanding HCMV latency and reactivation that will lead to the development of novel therapies to prevent viral reactivation in vulnerable patients.

## Methods

### Cells and Viruses

Details regarding the cells and their culture conditions, as well as details on the propagation of the TB40/E BAC-derived viruses are included in the SI Appendix, SI Methods.

### DNA, RNA, and Protein Analyses

Experimental approaches are included in the SI Appendix, SI Methods.

### Infection of Kasumi-3 and CD34^+^ Cells

Kasumi-3 cells were infected as described previously (22, 26, 30). Briefly, cells were serum-starved in X-VIVO15 (Lonza) 48 hours (h) before infection and then infected at a multiplicity of 1.0 50% tissue culture infectious dose (TCID_50_)/cell by centrifugal enhancement (1,000 × *g*, 35 minutes [min], room temperature) at 5 × 10^5^ cells/ml in X-VIVO15. The following day, cells were treated with trypsin to remove any virus that had not entered the cell and then cushioned onto Ficoll-Pacque (GE Healthcare Life Sciences) to remove residual virus and debris. Infected cells were washed three times with phosphate-buffered saline (PBS), replated in X-VIVO15 at 5 × 10^5^ cells/ml, and harvested as indicated in the text. Reactivation in Kasumi-3 cells was induced with 20 nM 12-O-tetradecanoylphorbol-13-acetate (TPA; Fisher) for 48 h in X-VIVO15, while latently infected counterpart cultures were treated with equal volumes of dimethylsulfoxide (DMSO; vehicle) in X-VIVO15.

Isolation of CD34^+^ HPCs is described in detail elsewhere (27). Immediately following isolation, CD34^+^ HPCs were infected at a multiplicity of 2.0 TCID_50_/cell, as previously described (22, 26, 30, 46), in infection media consisting of IMDM supplemented with 10% BIT9500 serum substitute (Stem Cell Technologies), 2 mM L-glutamine, 20 ng/mL low-density lipoproteins, and 50 μM 2-mercaptoethanol. The next day, cultures were washed three times in PBS and replated in 0.4 μm-pore transwells (Corning) over irradiated murine stromal cells in human Long Term Culture Medium (hLTCM, MyeloCult H5100 [Stem Cell Technologies] supplemented with 1 μM hydrocortisone, and 100 U/ml each of penicillin and streptomycin), described in detail in the SI Appendix, SI Materials and Methods.

### Multistep Growth Analyses

Experimental details are included in the SI Appendix, SI Methods.

### Extreme Limiting Dilution Assay

Reactivation efficiency in Kasumi-3 cells or CD34^+^ HPCs was measured by extreme limiting dilution assay (ELDA), as described previously (27). Briefly, Kasumi-3 cells were latently infected for 7 d and then co-cultured with naïve NuFF-1 cells in the presence either of vehicle (DMSO) or 20 nM TPA to maintain latency or induce reactivation, respectively. Infected cells were serially diluted two-fold onto naïve NuFF-1 cells and cultured for 14 d (as described in (27)). For CD34^+^ HPCs, cells were co-cultured with naïve NuFF-1 cells in a two-fold serial dilution as above, in the presence of hLTCM to maintain latency or reactivation media (RPMI supplemented with 20% FBS, 10 mM HEPES, 1 mM sodium pyruvate, 2 mM _L_-glutamate, 0.1 mM 0.1 mM non-essential amino acids, 100 U/ml each of penicillin and streptomycin with 15 ng/ml each of IL-6, G-CSF, GM-CSF and IL-3 [all from R&D Systems]). The production of infectious virus was quantified using viral-expressed mCherry as a marker of infection and ELDA software (bioinf.wehi.edu.au/software/elda/index.html).

### Chromatin Immunoprecipitation

Experimental details are provided in the SI Appendix, SI Methods.

## Acknowledgments

This work was supported by the American Heart Association Scientist Development Grant 15SDG23000029 (to C.M.O’C.) and Cleveland Clinic funding (to C.M.O’C.).

