## Supplemental Information for "Activator protein-1 transactivation of the major immediate early locus is a determinant of cytomegalovirus reactivation from latency"

**This PDF file includes:**

Supplementary text
Figures S1 to S8
Table S1
SI References

**Other supplementary materials for this manuscript include the following:**

n/a

### Supplementary Information Text

### SI Methods

#### Cells and Viruses

All cells were maintained at 37°C and 5% $CO_2$. NuFF-1 (primary newborn human fibroblasts, GlobalStem; passages 13 - 26) were maintained in Dulbecco's modified Eagle medium (DMEM), supplemented with 10% fetal bovine serum (FBS), 2 mM L-glutamine, 0.1 mM non-essential amino acids, 10 mM HEPES, and 100 U/ml each of penicillin and streptomycin. Retinal pigmented epithelial cells (ARPE-19; ATCC passages 34 - 38) were maintained in 1:1 DMEM-F12 medium containing 10% FBS, 2.5 mM L-glutamine, 0.5 mM sodium pyruvate, 15 mM HEPES, 1.2 g/L sodium bicarbonate, and 100 U/ml each of penicillin and streptomycin. Kasumi-3 cells (ATCC) were cultured in Roswell Park Memorial Institute (RPMI) 1640 medium (ATCC, catalog no. 30-2001) supplemented with 20% FBS, 100 U/ml each of penicillin and streptomycin, and 100 µg/ml gentamicin at a density of $5 \times 10^5$ – $1 \times 10^6$ cells/ml. Primary $CD34^+$ HPCs were isolated from de-identified cord blood samples (Abraham J. & Phyllis Katz Cord Blood Foundation *d.b.a.* Cleveland Cord Blood Center & Volunteer Donating Communities in Cleveland and Atlanta) by magnetic separation, as described elsewhere (1). Murine stromal cells, M2-10B4 (MG3) and S1/S1, were gifts from Terry Fox Laboratories, BC Cancer Agency, Vancouver, BC, Canada. MG3 cells were propagated in RPMI 1640, supplemented with 10% FBS and 100 U/ml each of penicillin and streptomycin. S1/S1 cells were propagated in Iscove's modified Dulbecco's medium (IMDM), supplemented with 10% FBS, 1 mM sodium pyruvate, and 100 U/ml each of penicillin and streptomycin. MG3 and S1/S1 cells were plated in a 1:1 ratio (~$1.5 \times 10^5$ cells of each cell type) onto collagen-coated (1 mg/ml) 6-well plates in human $CD34^+$ long-term culture media (hLTCM), consisting of MyeloCult H5100 (Stem Cell Technologies) supplemented with 1 µM hydrocortisone, and 100 U/ml each of penicillin and streptomycin. The following day, the cells were irradiated using a fixed source $^{137}$Cesium, Shepherd Mark I Irradiator at 20 Gy. These cells were used as feeder cells for the $CD34^+$ HPC infectivity assays described in the main text, Materials and Methods.

Where indicated, cells were treated with 20 nM T-5224 (BioVision) a concentration we previously showed does not impact cell viability (2), to inhibit c-fos and AP-1 signaling. Equal volumes of DMSO were used as a vehicle control.

Bacterial artificial chromosome (BAC)-derived TB40/E*mCherry* (WT) (3) was used to generate the AP-1 proximal binding site mutant, TB40/E*mCherry*-proximal-AP-*1mutant* (AP-1$mut_p$), and the AP-1 distal binding site mutant, TB40/E*mCherry*-distal-AP-1-*mutant* (AP-1$mut_d$) by *galK* recombineering, as previously described (4). Once verified by Sanger sequencing, AP-1$mut_p$ was used to repair the mutation to wild type sequence to generate TB40/E*mCherry*-proximal-AP-1*repair* (AP-1$rep_p$), while AP-1$mut_d$ was used to repair the mutation to wild type sequence to generate TB40/E*mCherry*-distal-AP-1*repair* (AP-1$rep_d$). To generate the recombinant virus in which both the AP-1 proximal and distal sites were mutated, AP-1$mut_{pd}$, we mutated the distal site in the AP-1$mut_p$ background. After verification by Sanger sequencing, AP-1$mut_{pd}$ was used to repair both AP-1 binding sites to their wild type sequence, AP-1$rep_{pd}$. The primers used for *galK* insertions, the double stranded oligonucleotides (ds oligo) used for the reversions, and the primers used to sequence verify the clones are listed in SI Appendix, Table S1.

Viral stocks were generated as described previously [e.g.: ref. (5)]. All viral stocks were titered by 50% tissue culture infectious dose ($TCID_{50}$) assay on NuFF-1 and ARPE-19 cells.

#### Multistep Growth Analyses

NuFF-1 or ARPE-19 cells were infected to evaluate multistep viral growth properties. NuFF-1 cells were infected at a multiplicity of 0.01 $TCID_{50}$/cell for 1 h with gentle rocking every 15 minutes (min), after which cells were washed twice in phosphate buffered saline (PBS) and returned to culture media. Infectious supernatants from each fibroblast culture were collected, and the cultures were replenished with fresh media at the time points indicated in the text. ARPE-19 cells were infected at a multiplicity of 0.1 $TCID_{50}$/cell using centrifugal enhancement (1000 x *g*, 30 min, room temperature), followed by 2 h incubation with gentle rocking every 15 min. Infected ARPE-19 cells were then washed twice in PBS and returned to culture media. ARPE-19 cells and media were collected at the indicated times post-infection, and cells were disrupted by three freeze-thaw cycles, after which cell debris was removed by centrifugation. To assess viral titers following infection of NuFF-1 and ARPE-19 cells, infectious virus was titrated on naïve NuFF-1 cells, and titers were determined by $TCID_{50}$ assay.

**DNA, RNA, and Protein Analyses**

Total RNA was isolated from cells using the High Pure RNA Isolation Kit (Roche), in accordance with the manufacturer's instructions. Next, cDNA (1 µg per sample) was reverse transcribed with the TaqMan RT reagent kit (Applied Biosystems) using random hexamers, as previously described (6). Transcripts were quantified using 20 ng (viral transcripts) or 4 ng (cellular transcript) of cDNA template for quantitative polymerase chain reaction (qPCR) using SYBR green PCR mix (Applied Biosystems). The absolute abundance of each transcript was calculated by comparison to a standard curve. BAC DNA, termed TB40/E-BACstandard, was used to generate transcript-specific standard curves, with the exception of the data in Fig 5 (see below). The TB40/E-BACstandard also contains fragments of cellular glyceraldehyde-3-phosphate dehydrogenase (GAPDH) and MDM2 within intergenic regions of the BAC. TB40/E-BACstandard DNA was purified and the concentration determined using a Nanodrop spectrophotometer. Ten-fold serial dilutions of the TB40/E-BACstandard were analyzed in duplicate for each viral specific transcript and the cellular *GAPDH* control. All experimental samples were analyzed in triplicate and then normalized to cellular *GAPDH*. All primers for RT-qPCR analysis are shown in SI Appendix, Table S1.

RNA isolation and reverse transcription for the data in Fig. 5 were performed as above. The abundance of each transcript was quantified by comparison to a standard curve specific to each primer pair, as described in detail previously (7). Prior to use, each primer pair was verified for: specificity (e.g. amplification of the specific MIE alternative promoter-derived sequence and not others), efficiency (>70% efficiency for each primer pair), and linear range (linear between $10^9$ and $10^4$; $R^2$ > 0.95). The primer pair used to detect dP (UTR487) as detailed by Arend et. al (6), were not suitable for AP-1$mut_p$, as the reverse primer targeted the promoter proximal AP-1 binding site, thus, a newly designed primer was utilized. Primers used herein are included in SI Appendix, Table S1.

For genome quantification, cells were washed three times in PBS and briefly treated with trypsin to remove any cell-bound virus that failed to enter. Cells were then washed and lysed by incubation at 37°C in lysis buffer (400 mM NaCl, 10 mM Tris, pH 7.5, 10 mM EDTA, 0.5% SDS, 100 µg/ml proteinase K) overnight, followed by RNAseA (100 µg/ml) treatment for 1 h at 37°C, extracted with phenol-chloroform-isoamyl alcohol, and precipitated with ethanol. DNA was resuspended in 10 mM Tris, pH 8.0 with 0.1 mM EDTA, and equal volumes were used in the qPCR reaction. The absolute amounts of cell-associated viral DNA were calculated by comparison to a standard curve, as described above, and then normalized to the cellular gene MDM2 (SI Appendix, Table S1).

**Chromatin Immunoprecipitation**

Cells were infected as indicated for 7 d and then treated with vehicle (DMSO) or 20 nM TPA for an additional 2 d. Cells were then fixed in 1% formaldehyde for 15 min at room temperature and then quenched with 125 mM glycine. Cells were lysed in immunoprecipitation (IP) buffer (150 mM NaCl, 50 mM Tris·HCl, pH 7.5, 5 mM EDTA, 0.5% Igepal-CA630, 1% Triton X-100) and debris was removed by centrifugation. DNA was sheared to 0.3- to 1-kb fragments with a MiSonix

Sonicator 3000 (20% output, 0.5 seconds on/off, 1 min) and aliquots stored as input controls. DNA associated with AP-1 was immunoprecipitated with protein A agarose (Millipore Sigma) and anti-fos (Cell Signaling #2250, diluted 1:100) or anti-jun (Cell Signaling #9165, diluted 1:50) antibodies using normal rabbit serum (Cell Signaling) as a control for non-specific binding. DNA was eluted by boiling and was followed by proteinase K treatment (0.2 ng/µl). DNA from disrupted nucleosomes was precipitated and used in PCR with primers directed at the MIEP or the UL69 non-promoter region (SI Appendix, Table S1). To analyze fos or jun binding to the MIE enhancer, we used primers spanning an 150 bp region. These primers are located between -30 to -9 (forward) and +97 to +119 (reverse), relative to the +1 site. Note that the distal site is located between +231 and +239 and the proximal site between +168 and +174 (relative to the +1 site), rendering their distance too close to evaluate separately.

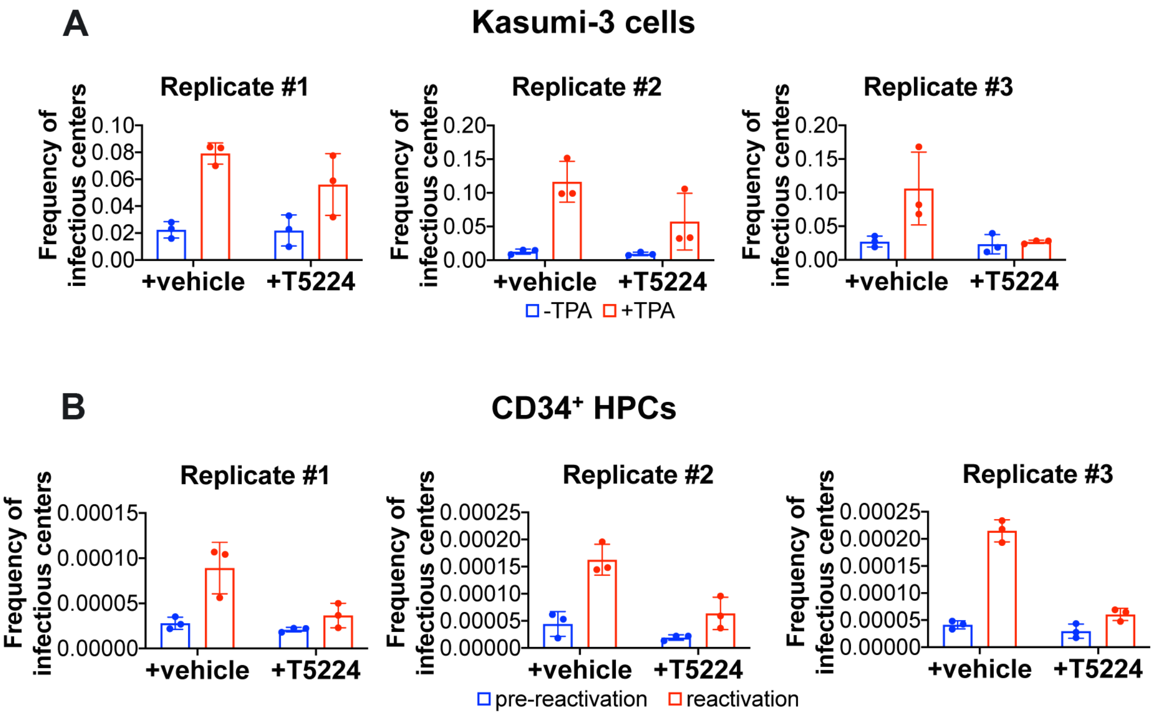


**Fig. S1. Biological replicates for the ELDA data presented in Fig1. (A)** Kasumi-3 cells (moi = 1.0) or **(B)** CD34$^+$ HPCs (moi = 2.0) were infected for 7 d under latent conditions with WT. The frequency of infectious centers was quantified by Extreme Limiting Dilution Analysis (ELDA) with **(A)** DMSO (-TPA, blue bars) or with TPA (+TPA, red bars) or **(B)** hLTCM (pre-reactivation, blue bars) or reactivation media (red bars), in the presence of vehicle (DMSO) or the c-fos inhibitor, T5224 (20 nM). Each data point (circles) represents an individual technical replicate.

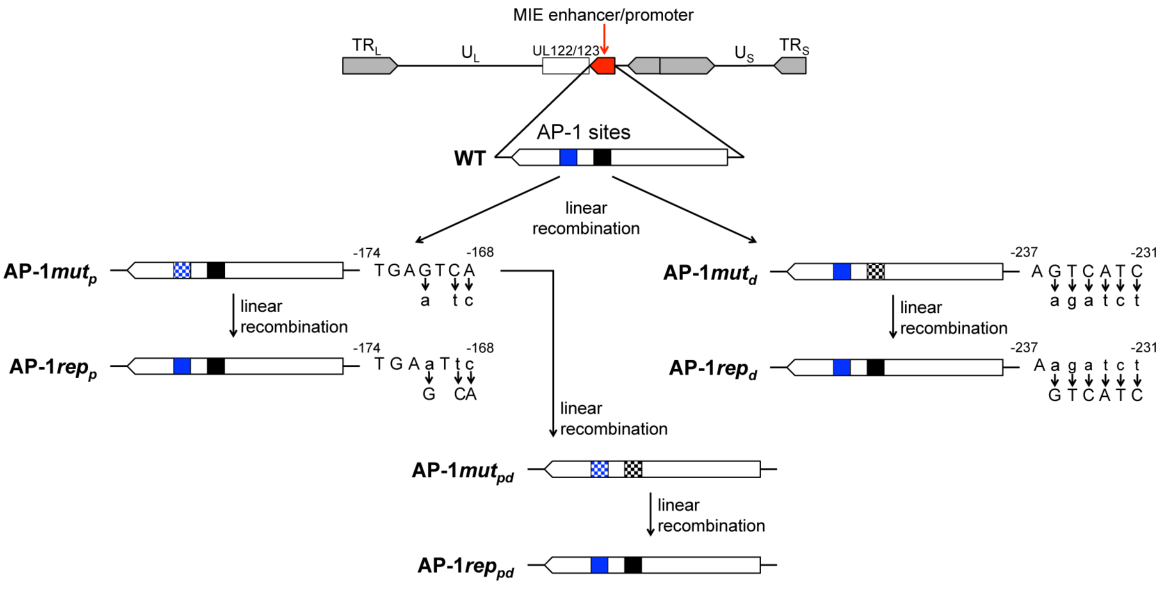


**Fig. S2. Schematic depicting the generation of viral recombinants.** BAC-derived TB40/E*mCherry* (WT) was used to generate: 1) a mutation in the proximal AP-1 binding site (blue box) of the MIE enhancer/promoter region (red), TB40/E*mCherry*-proximal-AP-1*mutant* (AP-1$mut_p$), or 2) a mutation in the distal AP-1 binding site (black box), TB40/E*mCherry*-distal-AP-1*mutant* (AP-1$mut_d$). The altered sequence for each single mutant is depicted at right, wherein the WT sequence is shown on the top (capital letters), with arrows corresponding to the altered bases (lower case letters). The position of the sequence, relative to the transcription start site, is shown above the WT sequence in each case. To return each mutated site to its wild type sequence: 1) AP-1$mut_p$ was used to repair the altered sequence to generate TB40/E*mCherry*-proximal-AP-1*repair* (AP-1$rep_p$) or 2) AP-1$mut_d$ was used to repair the altered sequence to generate TB40/E*mCherry*-distal-AP-1*repair* (AP-1$rep_d$). Using the AP-1$mut_p$ virus, we generated a double mutant, AP-1$mut_{pd}$, in which both AP-1 binding sites are mutated. Both mutated AP-1 binding sites in the AP-1$mut_{pd}$ double mutant were then repaired to their wild type sequences, yielding AP-1$rep_{pd}$. Solid blue box, wild type promoter proximal AP-1 site; solid black box, wild type promoter distal AP-1 site; blue/white checkered box, mutated promoter proximal AP-1 binding site; black/white checkered box represents, mutated promoter distal AP-1 binding site.

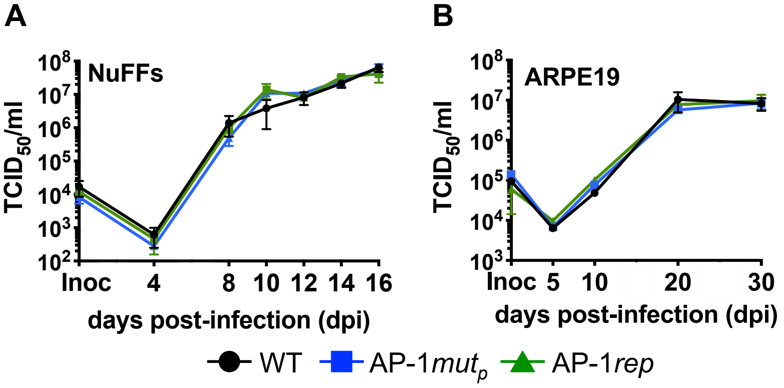


**Fig. S3. AP-*1mut*$_p$ grows to WT titers in lytically infected fibroblast and epithelial cells.** Multistep growth analyses were performed in **(A)** NuFF-1 (multiplicity of infection [moi] = 0.01) or **(B)** ARPE-19 cells (moi = 0.1). **(A)** Viral supernatants or **(B)** cell-associated virus were collected at the indicated dpi. **(A,B)** Viral titers were quantified by $TCID_{50}$ following infection of naïve NuFF-1 cells. Samples were analyzed in triplicate. Error bars represent the standard deviation of the technical replicates. A representative growth curve is shown for each cell type (n = 3). Unpaired t-test analyses with Benjamini, Krieger, Yekutieli false discovery rate approach revealed the data was not statistical significant at any time points evaluated.

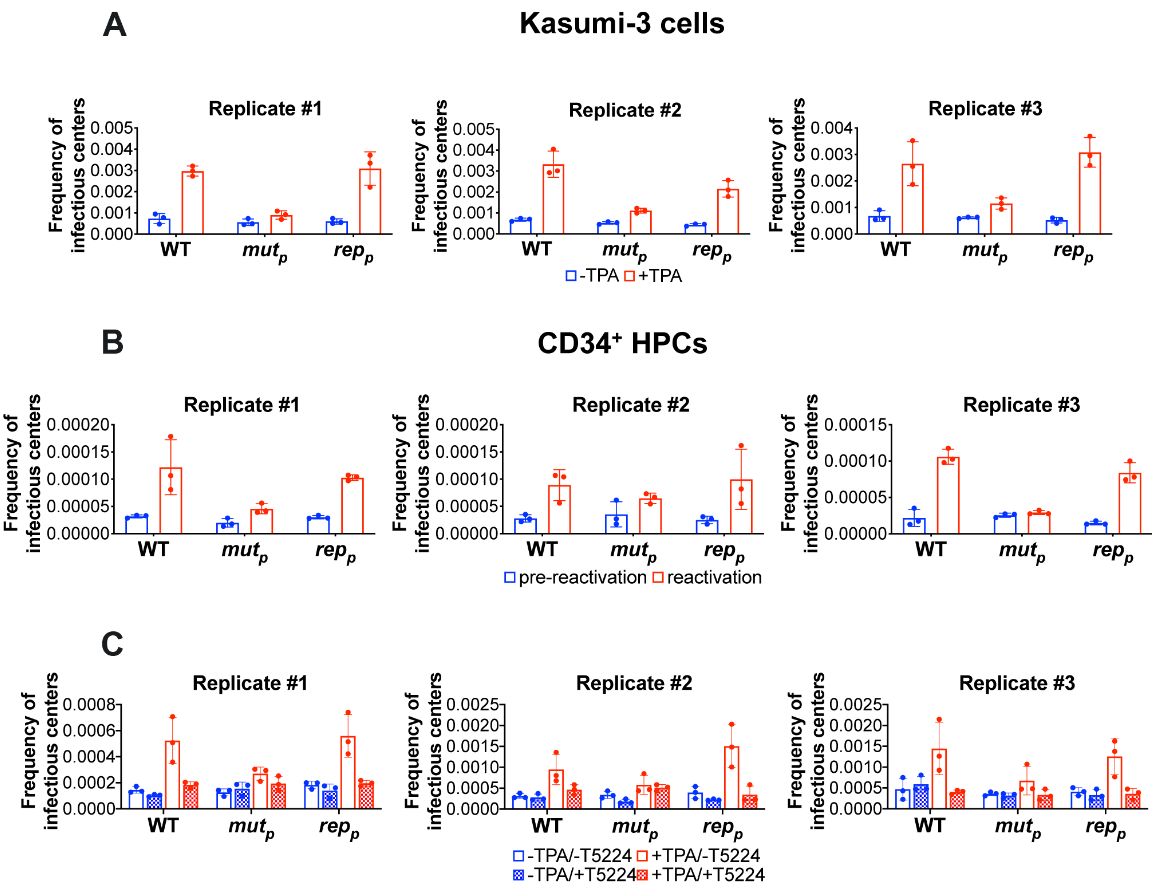


**Fig. S4. Biological replicates for the ELDA data presented in Fig2. (A,C)** Kasumi-3 cells (moi = 1.0) or **(B)** cord blood-derived CD34$^+$ HPCs (moi = 2.0) were infected with the indicated viruses. At 7 dpi, half of each infected population was cultured for an additional 2 d **(A,C)** with vehicle (DMSO; -TPA, blue bars) or TPA (+TPA, red bars) or **(B)** with hLTCM (pre-reactivation, blue bars) or reactivation (red bars) media. **(A,B)** Cells were then co-cultured with naïve NuFF-1 cells to quantify the frequency of infectious centers by ELDA. **(C)** Cells were co-cultured with naïve NuFF-1 cells in the presence of vehicle (DMSO; -T5224, open bars) or the fos inhibitor, T5224 (checked bars), and the frequency of infectious centers was quantified by ELDA. **(A-C)** Each data point (circles) represents an individual technical replicate. AP-1$mut_p$ ($mut_p$), AP-1$rep_p$ ($rep_p$).

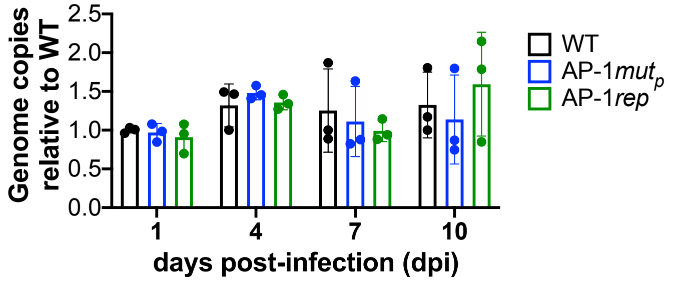


**Fig. S5. AP-1*mut*$_p$-infected Kasumi-3 cells maintain the viral genome during latent infection.** Kasumi-3 cells were infected (moi = 1.0) with the indicated viruses over a 10 d time course in conditions favoring latency. At the indicated dpi, cells were harvested, and cell-associated viral genomes were quantified by qPCR. Viral genome copies are shown relative to WT-infected cells (black bars) on day 1. Each data point (solid circles) is the mean of 3 technical replicates. Error bars indicate standard deviation of three biological replicates. The statistical significance was calculated using two-way ANOVA followed by Tukey's post-hoc analysis. Results were not statically significant.

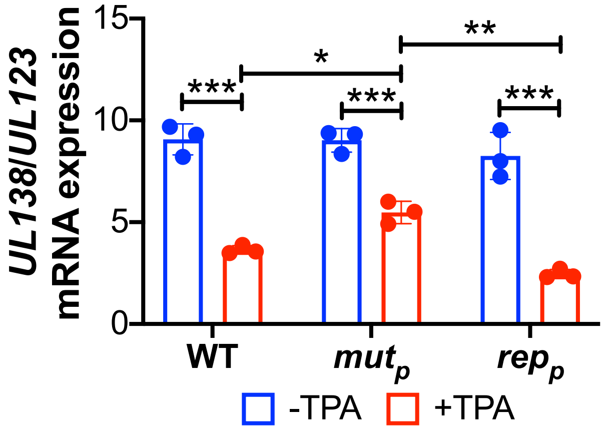


**Fig. S6. AP-1*mut*$_p$-infected Kasumi-3 cells express high levels of the latency-associated gene *UL138*.** Kasumi-3 cells were infected (moi = 1.0) with the indicated viruses over a 7 d time course in conditions favoring latency. At 7 dpi, half of each infected population was cultured for an additional 2 d with vehicle (DMSO; -TPA, blue bars) or TPA (+TPA, red bars). *UL138* and *UL123* mRNA expression were then quantified by RT-qPCR, plotted as a ratio of *UL138*/*UL123* and normalized to cellular *GAPDH*. Each data point (circles) is the mean of 3 technical replicates. Error bars indicate standard deviation of three biological replicates, and the statistical significance was calculated using two-way ANOVA followed by Tukey's post-hoc analysis. *$p<0.05$, **$p<0.01$, ***$p<0.001$; abbreviations: AP-1*mut*$_p$ (*mut*$_p$), AP-1*rep*$_p$ (*rep*$_p$).

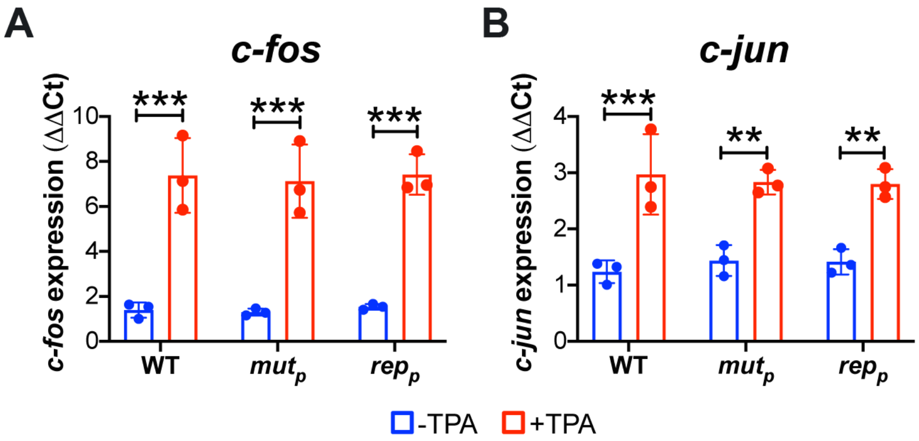


**Fig. S7. Transcription of *c-fos* and *c-jun* is not impacted by AP-1*mut*$_p$ infection and is increased in all virally infected cells following TPA treatment.** Kasumi-3 cells were infected (moi = 1.0) with the indicated viruses over a 7 d time course in conditions favoring latency. At 7 dpi, half of each infected population was cultured for an additional 2 d with vehicle (DMSO; -TPA, blue bars) or TPA (+TPA, red bars). **(A)** ***C****-fos* and **(B)** *c-jun* mRNA expression were then quantified by RT-qPCR. Each data point (circles) is the mean of three technical replicates. Error bars indicate standard deviation of three biological replicates, and the statistical significance was calculated using two-way ANOVA followed by Tukey's post-hoc analysis. **$p$<0.01, ***$p$<0.001. Abbreviations: AP-1*mut*$_p$ (*mut*$_p$), AP-1*rep*$_p$ (*rep*$_p$).

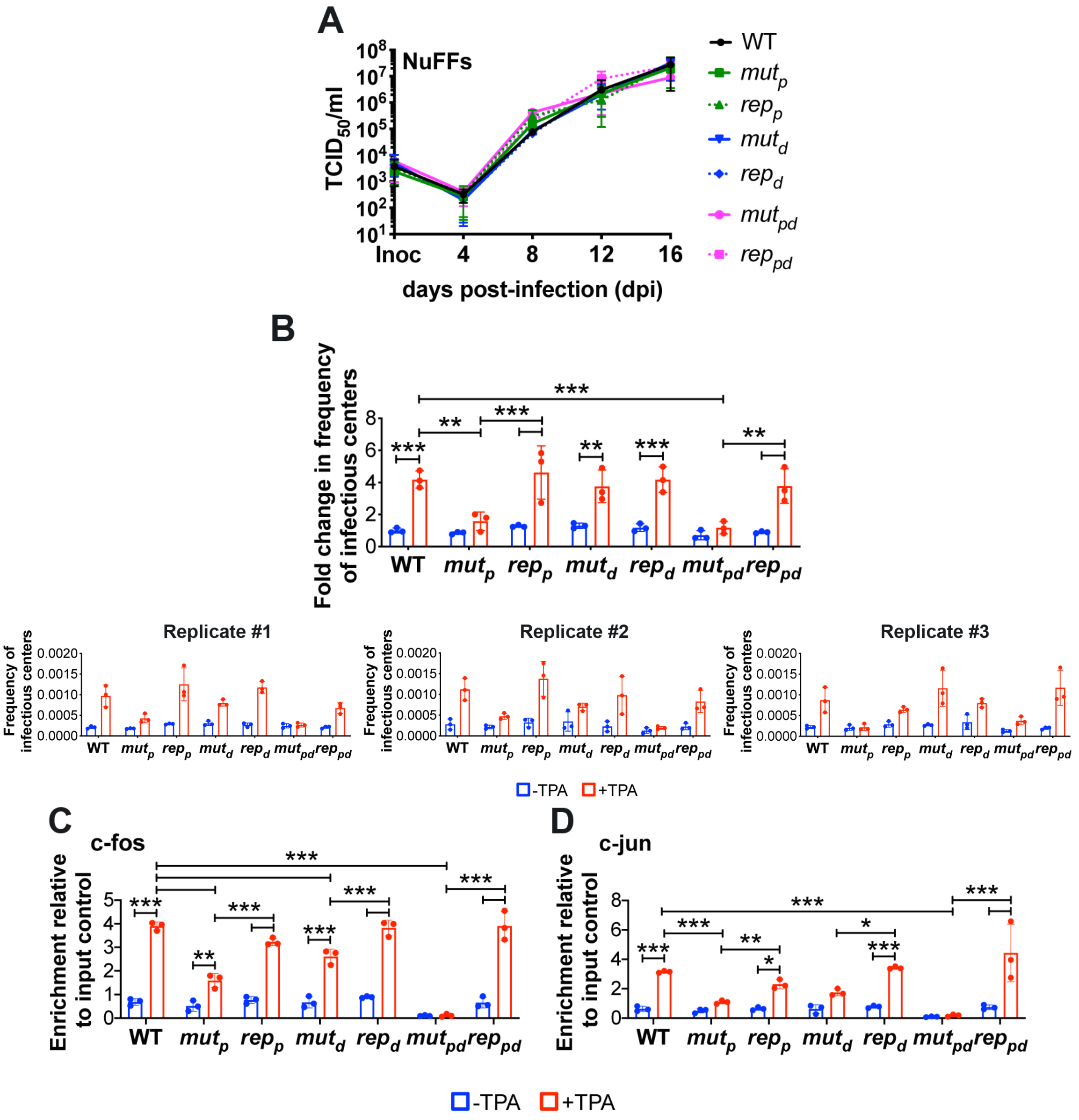


**Fig. S8. Mutation of the AP-1 promoter distal binding site does not significantly impact viral reactivation. (A)** Multistep growth analysis were performed in NuFF-1 cells (moi = 0.01). Viral supernatants were collected at the indicated dpi, and viral titers were quantified by $TCID_{50}$ following infection of naïve NuFF-1 cells. Samples were analyzed in triplicate. Error bars represent the standard deviation of the technical replicates. A representative growth curve is shown (n = 3). Unpaired t-test analyses with Benjamini, Krieger, Yekutieli false discovery rate approach revealed the data was not statistical significant at any time points evaluated. **(B-D)** Kasumi-3 cells were infected (moi = 1.0) with the indicated viruses over a 7 d time course in conditions favoring latency. At 7 dpi, half of each infected population was cultured for an additional 2 d with vehicle (DMSO; -TPA, blue bars) or TPA (+TPA, red bars). **(B)** The frequency of infectious centers was determined by ELDA. **(B, top)** The fold-change in frequency of infectious centers is shown relative to WT -TPA. Each data point (circles) is the mean of three technical replicates. Error bars indicate standard deviation of three biological replicates (shown below). **(B, bottom)** Three biological replicates, labeled Replicate #1-3, are shown, in which each data point (circles) represents an individual technical replicate (n=3). **(C,D)** AP-1 transcription factor binding to the MIEP was quantified by ChIP using an **(C)** α-fos or **(D)** α-jun antibody. **(C,D)**

Co-precipitated MIEP was quantified by qPCR, and data are shown as fold change relative to input for WT -TPA. Each data point (circles) is the mean of three technical replicates. Error bars indicate standard deviation of three biological replicates. **(B-D)** Statistical significance was calculated using two-way ANOVA followed by Tukey's post-hoc analysis. *$p<0.05$, **$p<0.01$, ***$p<0.001$. Abbreviations: AP-1$mut_p$ ($mut_p$), AP-1$rep_p$ ($rep_p$), AP-1$mut_d$ ($mut_d$), AP-1$rep_d$ ($rep_d$), AP-1$mut_{pd}$ ($mut_{pd}$), AP-1$rep_{pd}$ ($rep_{pd}$).

**Table S1. Oligonucleotides used in this study.**

| Primer Use | Sequence¶ | Target |
|---|---|---|
| *galK* insertion (proximal mutant generation) | GCCCATTTGCGTCAATGGGGCGGAGTTATTACGACATTTT<br>GGAAAGTCCCCCTGTTGACAATTAATCATCGGCA | AP-1$mut_p$ fwr |
| | GACATGAGCCAATATAAATGTACACATTATGATATAGATGC<br>AACGTATGCTCAGCACTGTCCTGCTCCTT | AP-1$mut_p$ rev |
| ds oligo (proximal mutant generation) | CTCCAGGCGATCTGACGGTTCACTAAACGAGCTCTGCTTA<br>TATAGACCTCCCATAGTACACGCCTACCGCCCATTTGCGT<br>CAATGGGGCGGAGTTATTACGACATTTTGGAAAGTCCCGT<br>TGAATTTGGTGCCAAAACAAACTCCCATTGACGTCAATGG<br>GGTGGAGACTTGGAAATCCCCG**TGAaTtc**AACCGCTATCCA<br>CGCCCATTGATGTACTGCCAAAACCGCATCACCATGGTAA<br>TAGCGATGACTAATACGTAGATGTACTGCCAAGTAGGAAA<br>GTCCCGTAAGGTCATGTACTGGGCATAATGCCAGGCGGG<br>CCATTTACCGTCATTGACGTCAATAGGGGGCGTACTTGGC<br>ATATGATACACTTGATGTACTGCCAAGTGGGCAGTTTACC<br>GTAAATACTCCTCCCATTGACGTCAATGGAAAGTCCCTATT<br>GGCGTTACTATGGGAACCCACGTCATTATTGACGTCAATG<br>GGCGGGGGTCGTTGGGCGGTCAGCCAGGCGGGCCATTT<br>ACCGTAAGTTATGTAACGCGGAACTCCATATATGGGCTAT<br>GAACTAATGACCCCGTAATTGATTACTATTAATAACTAGTC<br>AATAATCAATGTCAACATGGCGGTCATATTGGACATGAGC<br>CAATATAAATGTACACATTATGATATAGATGCAACGTATGC<br>AATGGCCATTAGCCAATATTGATTTACGCTATATAACCAAT<br>GACTAATATGGCTAATGGCCAATAT | AP-1$mut_p$ ds oligo |
| *galK* insertion (proximal mutant repair) | GCCCATTTGCGTCAATGGGGCGGAGTTATTACGACATTTT<br>GGAAAGTCCCCCTGTTGACAATTAATCATCGGCA | AP-1$rep_p$ fwr |
| | GACATGAGCCAATATAAATGTACACATTATGATATAGATGC<br>AACGTATGCTCAGCACTGTCCTGCTCCTT | AP-1$rep_p$ rev |
| ds oligo (proximal mutant repair) | CTCCAGGCGATCTGACGGTTCACTAAACGAGCTCTGCTTA<br>TATAGACCTCCCATAGTACACGCCTACCGCCCATTTGCGT<br>CAATGGGGCGGAGTTATTACGACATTTTGGAAAGTCCCGT<br>TGAATTTGGTGCCAAAACAAACTCCCATTGACGTCAATGG<br>GGTGGAGACTTGGAAATCCCCG**TGAgTca**AACCGCTATCC<br>ACGCCCATTGATGTACTGCCAAAACCGCATCACCATGGTA<br>ATAGCGATGACTAATACGTAGATGTACTGCCAAGTAGGAA<br>AGTCCCGTAAGGTCATGTACTGGGCATAATGCCAGGCGG<br>GCCATTTACCGTCATTGACGTCAATAGGGGGCGTACTTGG<br>CATATGATACACTTGATGTACTGCCAAGTGGGCAGTTTACC<br>GTAAATACTCCTCCCATTGACGTCAATGGAAAGTCCCTATT<br>GGCGTTACTATGGGAACCCACGTCATTATTGACGTCAATG<br>GGCGGGGGTCGTTGGGCGGTCAGCCAGGCGGGCCATTT<br>ACCGTAAGTTATGTAACGCGGAACTCCATATATGGGCTAT<br>GAACTAATGACCCCGTAATTGATTACTATTAATAACTAGTC<br>AATAATCAATGTCAACATGGCGGTCATATTGGACATGAGC<br>CAATATAAATGTACACATTATGATATAGATGCAACGTATGC<br>AATGGCCATTAGCCAATATTGATTTACGCTATATAACCAAT<br>GACTAATATGGCTAATGGCCAATAT | AP-1$rep_p$ ds oligo |
| *galK* insertion (distal mutant generation) | TATCCACGCCCATTGATGTACTGCCAAAACCGCATCACCA<br>TGGTAATAGCATGAGTCTGAAAGAAAAAACAC | AP-1$mut_d$ fwr |
| | ACGTAGATGTACTGCCAAGTAGGAAAGTCCCGTAAGGTCA<br>TGTACTGGGCTCAGCACTGTCCTGCTCCTTG | AP-1$mut_d$ rev |

| | | |
|---|---|---|
| ds oligo (distal mutant generation) | CAAACCGCTATCCACGCCCATTGATGTACTGCCAAAACCG CATCACCATGGTAATAGC**agatctT**AATACGTAGATGTACTG CCAAGTAGGAAAGTCCCGTAAGGTCATGTACTGGGCATAA TGC | AP-1$mut_d$ ds oligo |
| *galK* insertion (distal mutant repair) | TATCCACGCCCATTGATGTACTGCCAAAACCGCATCACCA TGGTAATAGCATGAGTCTGAAAGAAAAAACAC | AP-1$rep_d$ fwr |
| | ACGTAGATGTACTGCCAAGTAGGAAAGTCCCGTAAGGTCA TGTACTGGGCTCAGCACTGTCCTGCTCCTTG | AP-1$rep_d$ rev |
| ds oligo (distal mutant repair) | CAAACCGCTATCCACGCCCATTGATGTACTGCCAAAACCG CATCACCATGGTAATAGC**gatgacT**AATACGTAGATGTACTG CCAAGTAGGAAAGTCCCGTAAGGTCATGTACTGGGCATAA TGC | AP-1-$rep_d$ ds oligo |
| ds oligo (double mutant repair) | CAAAACAAACTCCCATTGACGTCAATGGGGTGGAGACTTG GAAATCCCCG**TGAgTca**AACCGCTATCCACGCCCATTGATG TACTGCCAAAACCGCATCACCATGGTAATAGC**gatgacT**AAT ACGTAGATGTACTGCCAAGTAGGAAAGTCCCGTAAGGTCA TGTACTGGG | AP-1-$rep_{pd}$ ds oligo |
| *UL123* qPCR | GCCTTCCCTAAGACCACCAAT | UL123 fwr |
| | ATTTTCTGGGCATAAGCCATAATC | UL123 rev |
| *UL122* qPCR | ATGGTTTTGCAGGCTTTGATG | UL122 fwr |
| | ACCTGCCCTTCACGATTCC | UL122 rev |
| *UL99* qPCR | GTGTCCCATTCCCGACTCG | UL99 fwr |
| | TTCACAACGTCCACCCACC | UL99 rev |
| *GAPDH* qPCR | ACCCACTCCTCCACCTTTGAC | GAPDH fwr |
| | CTGTTGCTGTAGCCAAATTCGT | GAPDH rev |
| *MDM2* qPCR | CCCCTTCCATCACATTGCA | MDM2 fwr |
| | AGTTTGGCTTTCTCAGAGATTTCC | MDM2 rev |
| *UL44* qPCR | TACAACAGCGTGTCGTGCTCCG | UL44 fwr |
| | GGCGTGAAAAACATGCGTATCAAC | UL44 rev |
| *UL36* qPCR | TCCAGACCATGGAGCTCATGA | UL36 fwr |
| | AGGAACTCTTTGCGTTCTGGC | UL36 rev |
| *UL37* qPCR | GACGAAGTCCGATGAGGAGGATG | UL37 fwr |
| | TGGGACACTGGGCGTTGTTG | UL37 rev |
| UL69 non-promoter region ChIP qPCR | CTCGTCGTGTGACAGCAGGATG | UL69 fwr |
| | GAACTACAGCAACTCAGCCGTTTGA | UL69 rev |
| MIE region ChIP qPCR | AACAGCGTGGATGGCGTCTCC | MIE internal fwr |
| | GGCACCAAAATCAACGGGACTTT | MIE internal rev |
| *c-fos* qPCR | CACTCCAAGCGGAGACAGAC | Fos fwr |
| | AGGTCATCAGGGATCTTGCAG | Fos rev |
| *c-jun* qPCR | TCGACATGGAGTCCCAGGA | Jun fwr |
| | GGCGATTCTCTCCAGCTTCC | Jun rev |
| cDNA fragment generation (reverse primer for iP2, MIEP, dP) | GGTCACGGGTGTCTCGGGCCGT | exon2-3 (8) |
| cDNA fragment generation (reverse primer for iP1) | CAAGGACGGTGACTGACTC | iP1 rev (9) |
| cDNA fragment generation (forward primer for iP2; paired with exon2-3 primer) | ACAGACTAACAGACTGTTCC | UTR70 out (8) |

| | | |
|---|---|---|
| cDNA fragment generation (forward primer for iP1, paired with iP1 rev primer) | CGCTGACGCATTTGGAAGAC | UTR378§ out (8) |
| cDNA fragment generation (forward primer for MIEP, paired with exon2-3 primer) | TGACCTCCATAGAAGACACC | UTR136§ out (8) |
| cDNA fragment generation (forward primer for dP, paired with exon2-3 primer) | AAGTACGCCCCCTATTGACG | UTR487§ out(8) |
| iP1-derived transcript (qPCR) | CTTAAGGCAGCGGCAGAA | iP1 fwr (9) |
| | CAAGGACGGTGACTGACTC | iP1 rev (9) |
| iP2-derived transcript (qPCR) | TAGCTGACAGACTAACAGAC | iP2 fwr (9) |
| | AGGACTCCATCGTGTCAAGG | iP2 rev (9) |
| MIEP-derived transcript (qPCR) | TTGACCTCCATAGAAGACAC | UTR136§ fwr (8) |
| | CCTTGACACGATGGAGTCCT | UTR136§ rev (8) |
| dP-derived transcript (qPCR) | CTGCCAAAACCGCATCACCATGG | UTR487 rev |
| | GCATTATGCCCAGTACATGACC | UTR487§ fwr (8) |

¶Primer sequences are shown 5' to 3' in orientation. Underlined sequences denote those corresponding to the pGalK plasmid, described elsewhere (10). Bolded sequences in the ds oligos denote the AP-1 binding site sequences. Lowercase nucleotides in the AP-1$mut_p$, AP-1$mut_d$, and AP-1$mut_{pd}$ ds oligos denote those that were mutated. UTR, untranslated region; fwr, forward primer; rev, reverse primer; ds oligo, double stranded oligonucleotide. §Nomenclature for primers from published work is maintained. UTR136, canonical MIEP; UTR70, iP2 internal promoter; UTR378, iP1 internal promoter; UTR487, distal promoter (dP).
